# Serum amyloid alpha 1-2 are not required for systemic inflammation in the 4T1 murine breast cancer model

**DOI:** 10.1101/2022.09.26.509617

**Authors:** Chenfeng He, Riyo Konishi, Ayano Harata, Yuki Nakamura, Rin Mizuno, Mayuko Yoda, Masakazu Toi, Kosuke Kawaguchi, Shinpei Kawaoka

## Abstract

Cancers induce the production of acute phase proteins such as serum amyloid alpha (SAA) in the liver and cause systemic inflammation. Despite the well-known coincidence of acute phase response and systemic inflammation, the direct roles of SAA proteins in systemic inflammation in the cancer context remains incompletely characterized, particularly in vivo. Here, we investigate the in vivo significance of SAA proteins in systemic inflammation in the 4T1 murine breast cancer model. 4T1 cancers elevate the expression of SAA1 and SAA2, the two major murine acute phase proteins in the liver. The elevation of *Saa1-2* correlates with the up-regulation of immune cell-related genes including neutrophil markers. To examine this correlation in detail, we generate mice that lack *Saa1-2* and investigate immune-cell phenotypes. RNA-seq experiments reveal that deletion of *Saa1-2* does not strongly affect 4T1-induced activation of immune cell-related genes in the liver and bone marrow. Flow cytometry experiments demonstrate the dispensable roles of SAA1-2 in cancer-dependent neutrophil infiltration to the liver. This study clarifies the negligible contribution of SAA1-2 proteins in systemic inflammation in the 4T1 breast cancer model.

## Introduction

Systemic inflammation is a major phenomenon caused by solid cancers (Allen et al., 2020; Baazim et al., 2022; Biswas and Acharyya, 2020; Hiam-Galvez et al., 2021). Advanced solid cancers induce the proliferation of particular immune cell types, expression of inflammatory cytokines, and migration of immune cells to particular organs such as the liver. These abnormalities are generally associated with a worse prognosis for cancer patients (Fearon et al., 2011; Petruzzelli and Wagner, 2016; Stephens et al., 2008; Zhang et al., 2021). Understanding how cancer cells modulate the host immune system is thus an important area of research for developing therapeutics that could mitigate cancer’s adverse effects on the host.

Acute phase response, traditionally known as the increased plasma concentration of the liver-derived secretory proteins (i.e., acute phase proteins) upon stimuli, is widely observed in various animal cancer models and cancer patients (Biswas and Acharyya, 2020; Gruys et al., 2005; Lai et al., 2021; Sack, 2018; Stephens et al., 2008; Wu et al., 2022). Such acute phase proteins include serum amyloid alpha (SAA) proteins (Sack, 2018). In the presence of stimuli such as infection and cancers, hepatocytes produce a massive amount of SAA proteins (Hojo et al., 2017; Lee et al., 2019; Sack, 2018). The plasma concentration of SAA proteins consequently elevates, modulating the immune system in various manners. In this regard, acute phase proteins are considered liver-derived amplifiers of systemic immune response to stimuli.

Previous studies have suggested that SAA proteins harbor an activity to modulate the activity of particular immune cell types (Sack, 2018). For example, human SAA proteins promote cytokine release of neutrophils (Furlaneto and Campa, 2000; Sack, 2018). Macrophages are also a target of SAA (Gaiser et al., 2021; Patel et al., 1998; Sun et al., 2015). It has been shown that SAA proteins induce muscle atrophy through Toll-like receptors (Hahn et al., 2020). It is of note that these findings are often based on in vitro and ex vivo experiments using exogenous SAA proteins (Sack, 2018). On the other hand, how endogenous SAA proteins contribute to systemic inflammation in vivo has been relatively less studied. Accordingly, our knowledge of the in vivo significance of SAA proteins is relatively limited. Insufficient understanding of the roles of SAA proteins in systemic inflammation in vivo is at least in part owing to the relatively small number of studies using genetic knockouts of *Saa* genes.

In the present study, we address the effects of genetic deletion of SAA1 and SAA2 (hereafter described as SAA1-2), the two major SAA proteins in mice, on systemic inflammation caused by 4T1 breast cancers. We find that 4T1 breast cancers fully activate the host immune system even in the absence of SAA1-2 proteins, suggesting the dispensable roles of SAA1-2 in cancer-induced systemic inflammation in this particular breast cancer model.

## Results

### 4T1 breast cancers elevate SAA1-2 in the liver

We previously demonstrated that transplantation of 4T1 breast cancer cells to BALB/c female mice strongly increased the expression of *Saa1-2* mRNAs in the liver (Hojo et al., 2017). We utilized the RNA-seq datasets we recently published (Mizuno et al., 2022) to confirm that *Saa1-2* mRNAs were induced in the livers of 4T1 breast cancer-bearing mice (Fig. 1A). We further validated this observation using reverse transcription quantitative PCR (RT-qPCR) detecting both *Saa1* and *Saa2* whose nucleotide sequences are 95% identical (Fig. 1B and Fig. S1A and Table S1). We also found that 4T1 breast cancers elevated hepatic SAA1-2 at the protein level (Fig. 1C). Together, we concluded that transplantation of 4T1 breast cancers enhanced acute phase response in the liver, which is in line with other cancer models (Lee et al., 2019).

**Figure 1:**
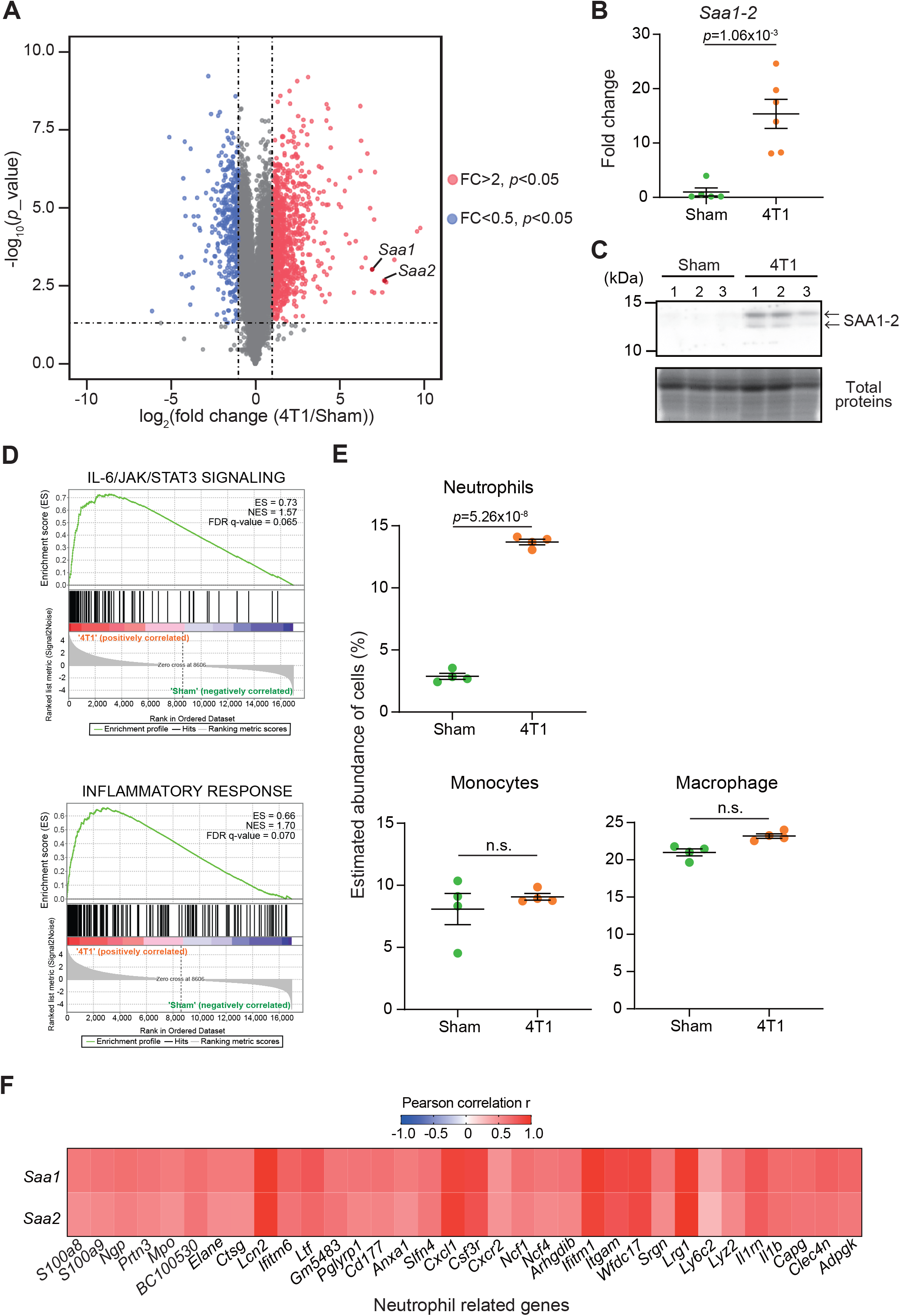
4T1 breast cancers elevate SAA1-2 in the liver. **(A)** RNA-seq experiments for the livers of sham-operated mice and 4T1-bearing mice in WT (14 days after 4T1 transplantation). A volcano plot (log_2_ fold average (4T1/sham) versus –log_10_ (*p* value)) of WT is shown. Genes showing more than 2-fold change with *p* < 0.05 are highlighted. *n* = 4. **(B)** qPCR analysis of *Saa1-2* in the livers of sham and 4T1-bearing mice. Averaged fold change data normalized to the sham group are presented as the mean ± SEM. The *p* value is shown (unpaired two-tailed Student’s *t*-test). *n* = 5 for sham-operated mice and *n* = 6 for 4T1-bearing mice. **(C)** Western blot analysis for SAA1-2 in the livers of sham and 4T1-bearing mice. *n* = 3. **(D)** GSEA plots that evaluate hepatic gene expression changes in “IL-6/JAK/STAT3 signaling” and “Inflammatory response” upon 4T1 transplantation. FDR *q* value, enrichment score (ES), and normalized enrichment score (NES) are shown. **(E)** Dot plots showing the estimated abundance of the indicated immune cell types in the liver of sham and 4T1-bearing mice. The scores are calculated using the RNA-seq datasets in Fig. 1A and ImmuCellAI-mouse. Data are mean ± SEM. The *p* value is shown (unpaired two-tailed Student’s *t*-test). n.s., not significant. **(F)** Heatmap representation of the correlations between *Saa1-2* and representative neutrophil-related genes at mRNA level using the RNA-seq dataset (A). *n* = 4. See also Fig. S1b.

To address whether the increased expression of SAA1-2 affects local and systemic inflammation, we characterized our liver transcriptome data using gene set enrichment analysis (GSEA) (Fig. 1D) (Mootha et al., 2003; Subramanian et al., 2005). Our analysis revealed that 4T1 breast cancer triggered various inflammatory responses in the livers as exemplified by the activation of the IL-6 signaling (Flint et al., 2016). This was in line with the previous observations that *Saa1-2* genes are under the control of the IL-6 signaling (Lee et al., 2019) and that solid cancers instigate the IL-6 signaling in the liver (Biswas and Acharyya, 2020; Flint et al., 2016). We then took advantage of ImmuCellAI-mouse to deduce the infiltration of various immune cells into the liver (Miao et al., 2021; Miao et al., 2020). Using this method, we found that neutrophils migrated into the liver upon cancer transplantation, which we and others previously reported (Fig. 1E) (Hojo et al., 2017; Lee et al., 2019). Furthermore, we noted the close correlations at the mRNA level between *Saa1-2* and other neutrophil-related genes such as *Lcn2* (Fig. 1F and Fig. S1B). Together with the known biochemical roles of SAA1-2 (Sack, 2018), these results led to a hypothesis that SAA1-2 proteins play some roles in immune cell activation in the presence of 4T1 breast cancers.

### Generation of mice completely lacking SAA1-2

To uncover the roles of SAA1-2 in inflammation in vivo in this model, we generated mice completely lacking *Saa1* and *Saa2*. These two genes are located closely in the murine genome (Fig. 2A), having redundant sequences and molecular functions (Fig. S1A) (Sack, 2018). We thus decided to delete the whole region encoding *Saa1* and *Saa2* genes by designing gRNAs on the right and left sides of this genomic locus (Fig. 2A) (Hojo et al., 2019). As a result, we succeeded in deleting both *Saa1* and *Saa2* as determined by genomic PCR (Fig. 2A-2B). Transplantation of 4T1 breast cancer cells to *Saa1-2* knockout mice no longer increased the expression of *Saa1-2* mRNAs (Fig. 2C) and proteins (Fig. 2D) in the liver. Western blot experiments demonstrated that two bands detected in a lysate prepared from the livers of 4T1-bearing mice disappeared in *Saa1-2* knockout mice (Fig. 1C and Fig. 2D). Adding recombinant SAA1 protein that lacks the signal peptide as a control, we reasoned that the lower band corresponds to the cleaved SAA proteins (Fig. S2). Taken together, we established mice where we completely canceled cancer-dependent increase in SAA1-2 proteins in the liver, allowing us to investigate the in vivo significance of these proteins in cancer-induced systemic inflammation.

**Figure 2:**
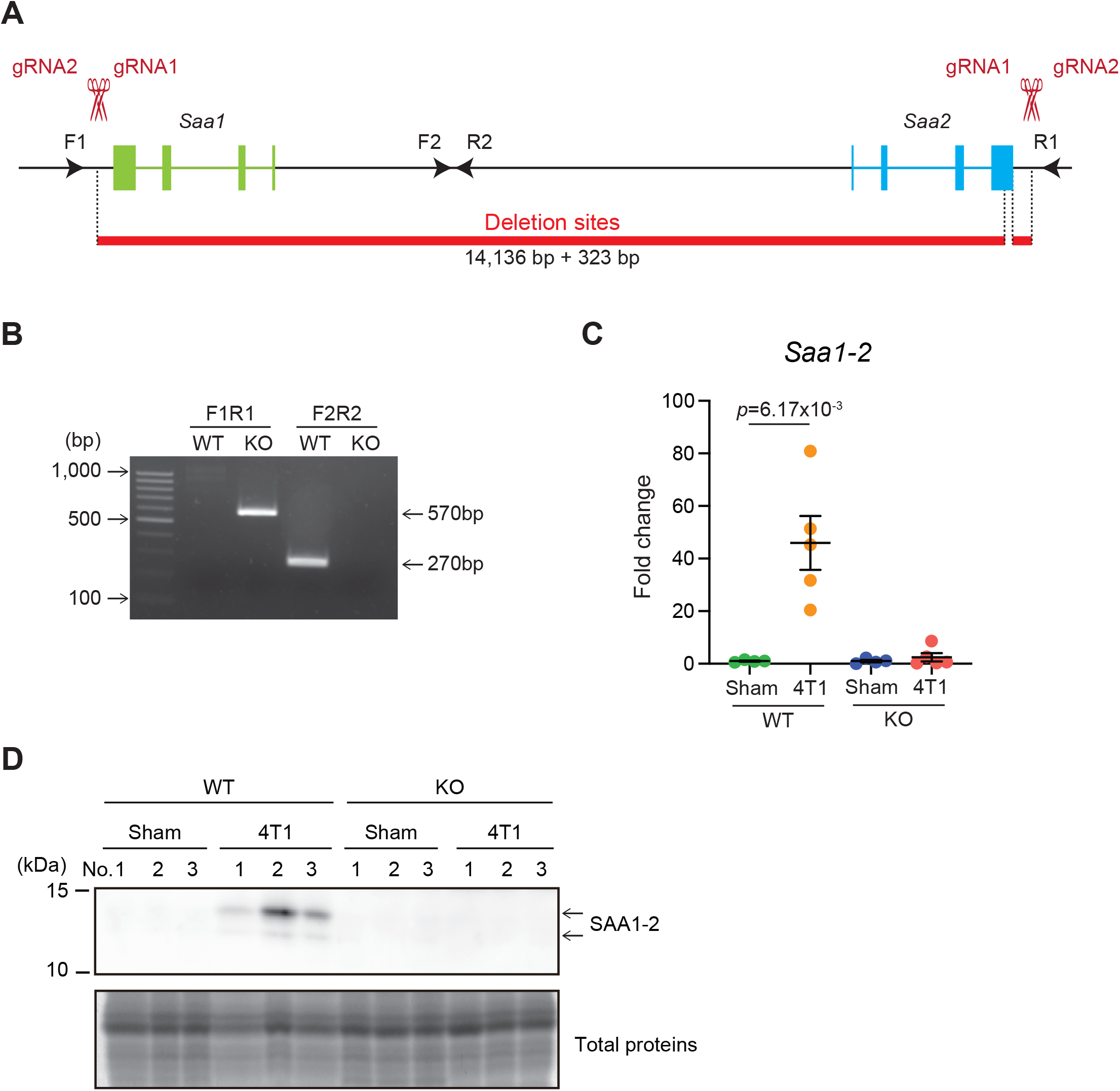
Generation of mice completely lacking SAA1-2. **(A)** Schematic representation of *Saa1-2* deletion using the CRISPR-Cas9 system. The gRNA-targeted sites and the primers used for genotyping experiments in (B) are indicated. The deleted regions are indicated in red. **(B)** A representative image of genomic PCR against the *Saa1-2* locus. **(C)** qPCR analysis of *Saa1-2* in the livers of sham and 4T1-bearing mice in WT and *Saa1-2* KO. Averaged fold change data normalized to the sham group in each genotype are presented as the mean ± SEM. The *p* value is shown (unpaired two-tailed Student’s *t*-test). *n* = 4 for sham-operated mice and *n* = 5 for the 4T1-bearing mice. **(D)** Western blot analysis for SAA1-2 in the livers of sham and 4T1-bearing mice in WT and *Saa1-2* KO. *n* = 3. See also Fig. S2.

### Deletion of *Saa1-2* does not have strong impacts on liver transcriptome

To thoroughly analyze the effects of *Saa1-2* knockout on the liver transcriptome, we performed RNA-seq analyses against the livers of WT and *Saa1-2* knockout mice (Fig. 3A and Table S2). We found that 4T1 breast cancer transplantation similarly affected liver transcriptome even in the absence of *Saa1-2* genes, as evidenced by the volcano plots shown in Fig. 3A. GSEA demonstrated that 4T1 breast cancers were capable of activating the IL-6 signaling and inflammatory response in the absence of *Saa1-2* genes (Fig. 3B), suggesting that the contribution of *Saa1-2* on liver transcriptome was subtle if any. In addition, we wanted to confirm these observations using different cohorts of 4T1 transplantation experiments. For this purpose, we quantified the mRNA expression of various immune cell marker genes in the liver, finding that *Saa1-2* KO did not have a significant impact on cancer-dependent up-regulation of representative immune-related genes in the liver (Fig. S3). These results suggested that 4T1 breast cancer cells do not require *Saa1-2* to increase the expression of immune-related genes in the liver in our experimental settings.

**Figure 3:**
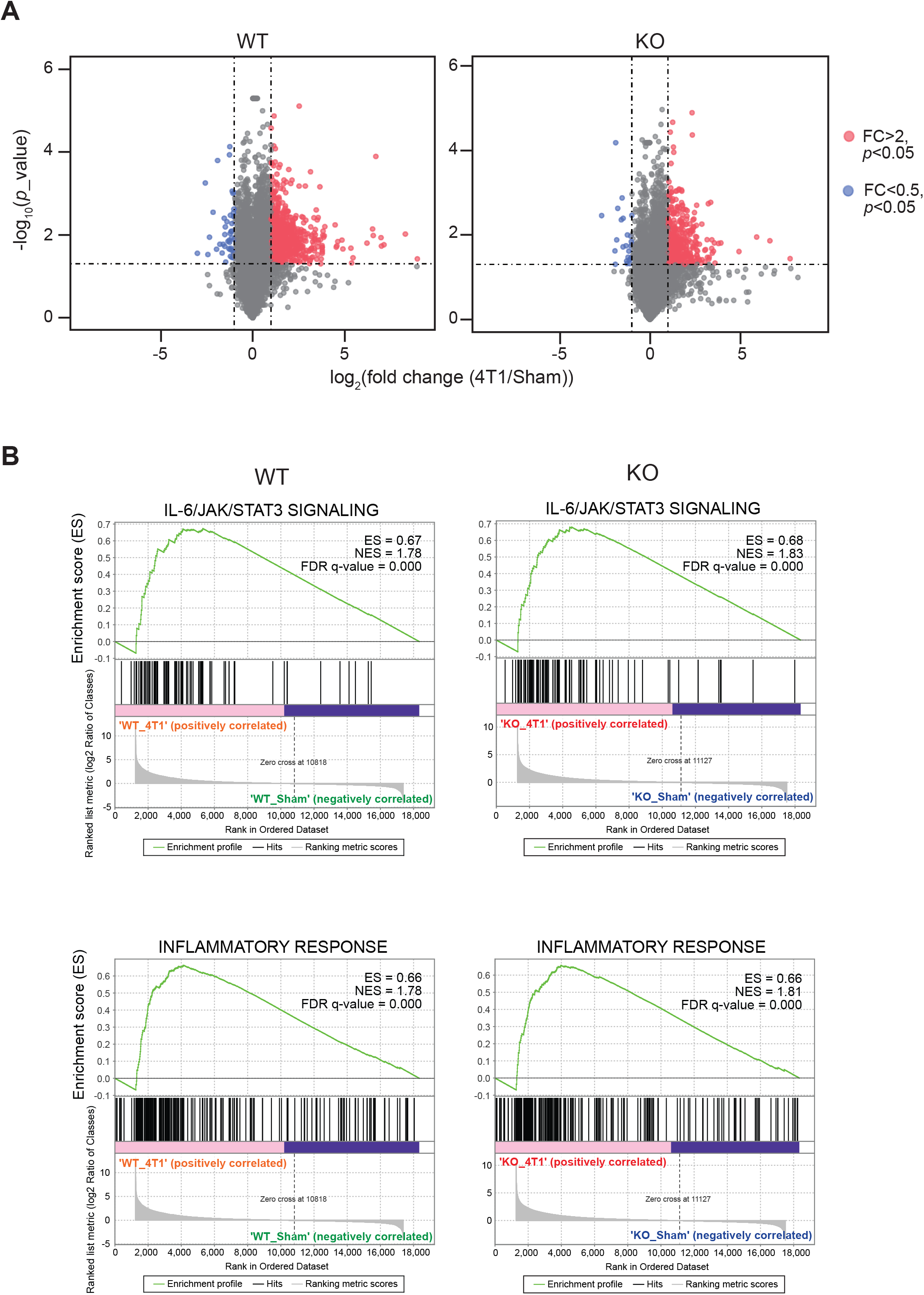
Deletion of *Saa1-2* does not have strong impacts on liver transcriptome. **(A)** RNA-seq experiments for the livers of sham-operated mice and 4T1-bearing mice in WT and *Saa1-2* KO (14 days after 4T1 transplantation). Volcano plots (log_2_ fold average (4T1/sham) versus –log_10_ (*p* value)) of WT (left) and *Saa1-2* KO (right) are shown. Genes showing more than 2-fold change with *p* < 0.05 are highlighted in red. *n* = 2 for the sham groups and *n* = 3 for 4T1-bearing groups. See also Fig. S3 for qPCR analyses using the different cohorts of 4T1 transplantation experiments. **(B)** GSEA plots that evaluate hepatic gene expression changes in “IL-6/JAK/STAT3 signaling” and “Inflammatory response” upon 4T1 transplantation in WT and *Saa1-2* KO. FDR *q* value, enrichment score (ES), and normalized enrichment score (NES) are shown.

### SAA1-2 are dispensable for 4T1-induced infiltration of immune cells to the liver

A previous study reported the significant contribution of SAA proteins in recruiting innate immune cells including neutrophils to the liver in the pancreatic cancer-bearing condition (Lee et al., 2019). This prompted us to further investigate the roles of SAA1-2 in immune cell recruitment to the liver in the 4T1 breast cancer model. To this end, we estimated the proportions of several immune cell types in the livers of cancer-bearing mice, comparing them between WT and *Saa1-2* KO.

ImmuCellAI-mouse analyses confirmed that transplantation of 4T1 breast cancer cells increased the proportion of neutrophils within the liver (Fig. 4A). Of note, the proportions of neutrophils in the liver were comparable between WT and *Saa1-2* KO in the cancer-bearing condition, implying negligible roles of SAA1-2 in recruiting neutrophils in this model. We also investigated the proportions of neutrophils, monocytes, and macrophages using flow cytometry. We collected immune cells from the livers of sham and cancer-bearing animals, quantifying those immune cells using a set of specific antibodies (see figure legends and methods). As shown in Fig. 4B, our data revealed that the proportions of neutrophils, monocytes, and macrophages were unaffected by *Saa1-2* KO. Collectively, although 4T1 breast cancer transplantation strongly induced SAA1-2 in the liver, it appeared that these acute phase proteins are not essential for cancer-dependent immune cell recruitments into the liver in the 4T1 model (Figs. 1-4).

**Figure 4:**
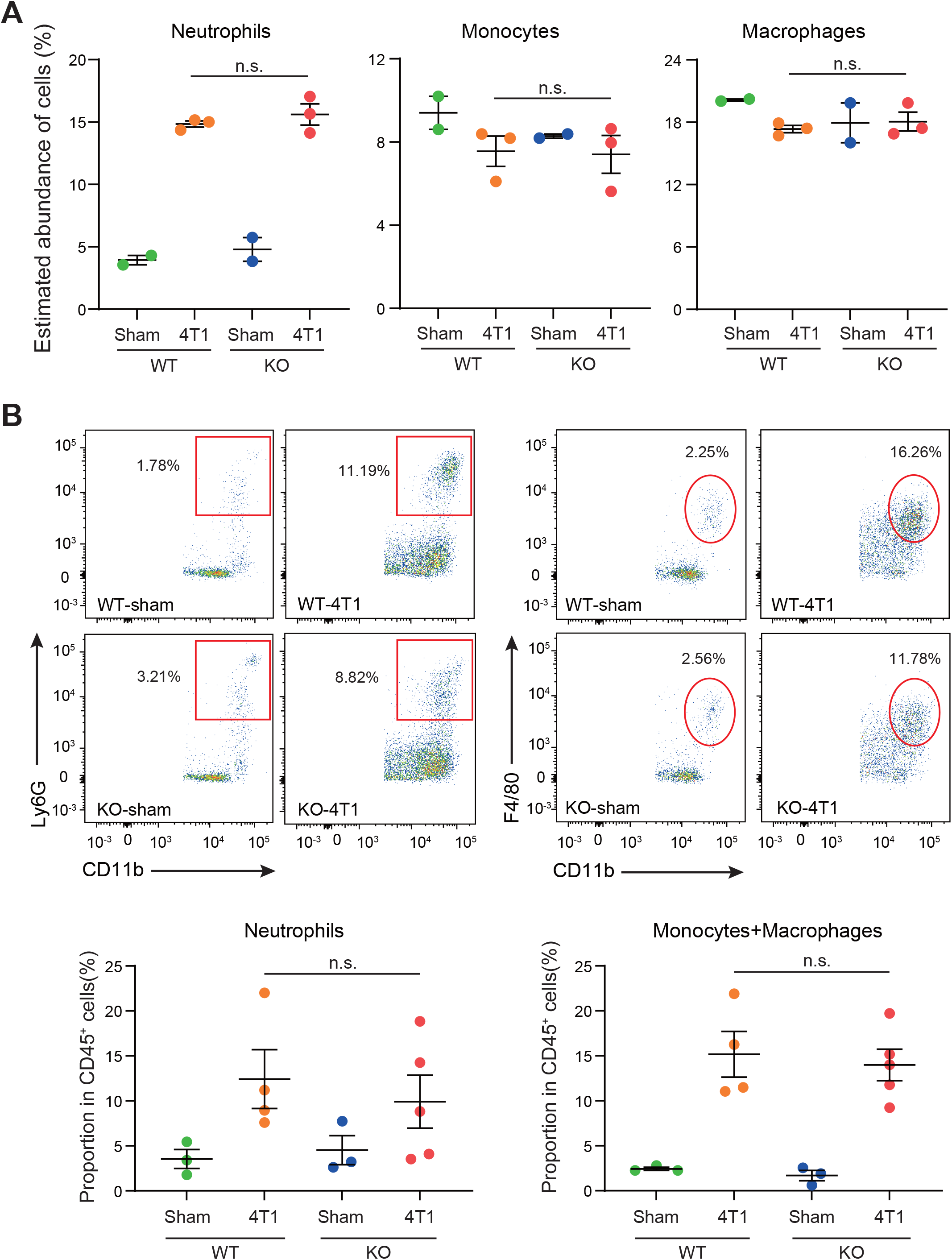
SAA1-2 are dispensable for 4T1-induced infiltration of immune cells to the liver. **(A)** Dot plots showing the estimated abundance of the indicated immune cell types in the liver of sham and 4T1-bearing mice. The scores are calculated using the RNA-seq dataset in Fig. 3A and ImmuCellAI-mouse. Data are presented as the mean ± SEM. n.s., not significant, unpaired two-tailed Student’s *t*-test. **(B)** Flow cytometric analysis of Ly6G^+^CD11b^+^ neutrophils and F4/80^+^CD11b^+^ monocytes and macrophages in the livers of sham and 4T1-bearing mice in WT and *Saa1-2* KO. Representative plots are shown. Data are mean ± SEM. The *p* value is shown (unpaired two-tailed Student’s *t*-test). *n.s*., not significant. *n* = 3 for the sham groups, *n* = 4 for 4T1-bearing WT mice, and *n* = 5 for 4T1-bearing *Saa1-2* KO mice. See also Fig. S4 for the gating strategies used in this study.

### Dispensable roles of SAA1-2 in 4T1-induced transcriptomic changes in the bone marrow

Innate immune cells such as neutrophils are born and matured in the bone marrow (Nauseef and Borregaard, 2014). It is also known that cancers affect immune cell development (Allen et al., 2020). Given these, it was likely that the altered immune cell status in the liver was owing to changes in the bone marrow. We thus wanted to extend our experiments on the roles of *Saa1-2* in immune cell activation in the bone marrow.

We performed RNA-seq experiments against immune cells collected from the bone marrow from WT and *Saa1-2* KO mice from which we obtained the liver transcriptome data (Fig. 3). We found that 4T1 transplantation affected gene expression in the bone marrow, resulting in many differentially expressed genes (Fig. 5A). According to GSEA, in WT, 4T1 transplantation activated the IL-6 signaling and inflammatory response (Fig. 5B), which is in line with our liver data (Fig. 3B). These results implied that inflammatory response observed in the liver is correlated with altered immune cell dynamics in the bone marrow. Moreover, ImmuCellAI-mouse demonstrated that 4T1 transplantation altered the proportions of various cell types including neutrophils in the bone marrow, supporting a cancer-induced reprogramming of the host immune system (Fig. 5C) (Allen et al., 2020). Most importantly, none of these immune cell phenotypes in the bone marrow was strongly buffered by the deletion of *Saa1-2*. We validated these data using qPCR with the different cohorts of experiments (Fig. S5). These results together provided evidence that the effects of *Saa1-2* KO on the bone marrow transcriptome were subtle if any, suggesting dispensable roles of SAA1-2 in 4T1-induced systemic reprogramming of the host immune system. Taken altogether, our results suggested that SAA1-2 proteins are not essential for local and systemic inflammation observed in the 4T1 breast cancer model.

**Figure 5:**
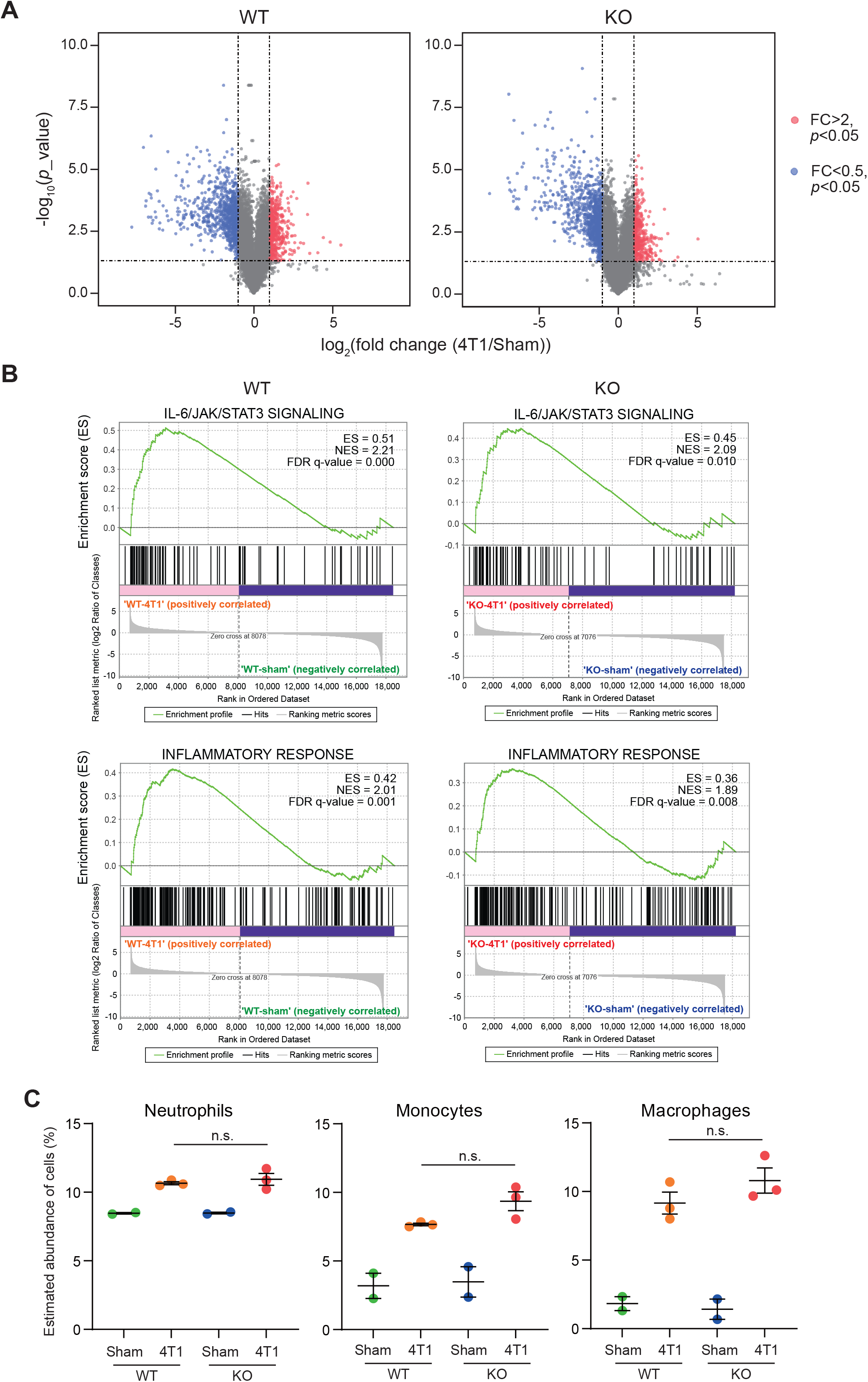
Dispensable roles of SAA1-2 in 4T1-induced transcriptomic changes in the bone marrow. **(A)** RNA-seq experiments for the bone marrows of sham-operated mice and 4T1-bearing mice in WT and *Saa1-2* KO (14 days after 4T1 transplantation). Volcano plots (log_2_ fold average (4T1/sham) versus –log_10_ (*p* value)) of WT (left) and *Saa1-2* KO (right) are shown. Genes showing more than 2-fold change with *p* < 0.05 are highlighted in red. *n* = 2 for the sham groups and *n* = 3 for 4T1-bearing groups. See also Fig. S5 for qPCR analyses using the different cohorts of 4T1 transplantation experiments. **(B)** GSEA plots that evaluate gene expression changes in the bone marrow in “IL-6/JAK/STAT3 signaling” and “Inflammatory response” upon 4T1 transplantation in WT and *Saa1-2* KO. FDR *q* value, enrichment score (ES), and normalized enrichment score (NES) are shown. **(C)** Dot plots showing the estimated abundance of the indicated immune cell types in the bone marrows of sham and 4T1-bearing mice. The scores are calculated using the RNA-seq dataset in Fig. 5A and ImmuCellAI-mouse. Data are presented as the mean ± SEM. The *p* value is shown (unpaired two-tailed Student’s *t*-test). n.s., not significant, unpaired two-tailed Student’s *t*-test.

## Discussion

Advanced cancers reprogram the host immune system, inducing systemic inflammation (Allen et al., 2020; Baazim et al., 2022; Biswas and Acharyya, 2020; Fearon et al., 2011; Hiam-Galvez et al., 2021; Stephens et al., 2008). Cancer-induced systemic inflammation is associated with the elevation in the levels of acute phase proteins both in murine cancer models and cancer patients, indicating the tight connection between systemic inflammation and acute phase response in the cancer contexts (Biswas and Acharyya, 2020; Gruys et al., 2005; Lai et al., 2021; Sack, 2018; Stephens et al., 2008; Wu et al., 2022). However, the causal relationship between them remains insufficiently addressed especially in vivo.

Using the 4T1 breast cancer model and *Saa1-2* KO mice, we evaluated the contribution of *Saa1-2* in 4T1-induced systemic inflammation. As shown in Fig. 1, the expression of *Saa1-2* showed significant correlations with immune cell gene expression and proportions of immune cells. In particular, the correlation between *Saa1-2* and innate immune cells including neutrophils appeared strong (Fig. 1). This prompted us to generate mice completely lacking both *Saa1* and *Saa2* (Fig. 2). Combining *Saa1-2* KO mice, RNA-seq, and flow cytometry, we found that, despite the strong correlation between SAA1-2 and various immune phenotypes, SAA1-2 proteins are not essential for the 4T1-induced immune cell alterations (Figs. 3-5). It thus appeared that 4T1 cancer cells could reprogram the host immune system independently of SAA1-2.

We do not exclude the possibility that our experimental settings had factors that mask the significance of SAA1-2. For example, the 4T1 breast cancer model induced the expression of *Saa3* (Fig. S6A). SAA3 is a protein whose amino acid sequence is approximately 62% similar to those of SAA1 and SAA2 (Fig. S6B). SAA3 plays important role in the pathogenesis caused by T helper 17 (Th17) cells (Lee et al., 2020). Our data demonstrated that *Saa3* was still induced in the liver and bone marrow of *Saa1-2* KO mice (Fig. S5 and S6A), possibly compensating for the absence of *Saa1-2*. It is also possible that 4T1-derived cytokines are sufficient to induce and maintain systemic inflammation (Kano, 2015). Given the massive number of transplanted 4T1 cancer cells in our experimental settings, the overwhelming amounts of cancer-derived cytokines might conceal the significance of SAA1-2 proteins produced by the liver. Our notion that experimental conditions might influence the roles of SAA1-2 does not contradict the previous report showing the critical role of SAA proteins in recruiting neutrophils to the liver in the presence of pancreatic cancers (Lee et al., 2019). In this regard, investigating SAA1-2 in other cancer models is critical to deepening our understanding of how SAA1-2 proteins contribute to specific pathophysiology in vivo. By doing so, we may reveal conditions critically affecting the in vivo significance of SAA1-2. Furthermore, SAA1-2 proteins may be important in phenomena we did not investigate in this study. For example, the inflammation status may have something to do with metabolism as we reported previously (Enya et al., 2018). The clarification of the in vivo functions of SAA1-2 in such phenomena awaits further examination.

In summary, we investigated the contribution of *Saa1-2* in systemic inflammation caused by 4T1 breast cancers, finding that *Saa1-2* genes are dispensable for systemic inflammation in this particular model. This study provides an example that the strong correlation in gene expression does not always mean the in vivo significance and deepens our understanding of the relationship between systemic inflammation and acute phase response in cancer contexts.

## Materials and Methods

### Mice

All animal experiment protocols were approved by the Animal Care and Use committee of Kyoto University. Mice were housed as described previously (Mizuno et al., 2022) in a 12-hour light/dark paradigm with food (CE-2, CLEA Japan, Inc., Tokyo, Japan) and water available *ad libitum*. Mice were randomly assigned to different experimental groups without any specific criterion. No blinding was performed. WT mice were purchased from Japan SLC Inc. (Hamamatsu, Japan).

### Generation of KO mice

BALB/c *Saa1-2* KO mice were generated as described previously (Hojo et al., 2019; Mizuno et al., 2022). In vitro fertilized eggs stored were thawed and electroporated using CUY-EDIT II (BEX, Tokyo, Japan) (amplitude 25V, duration 3 msec., interval 97 msec. for twice) with two independent gRNAs (25 ng/ml: FASMAC, Kanagawa, Japan), crRNAs (25 ng/ml: FASMAC), tracrRNA (25 ng/ml: FASMAC), and purified recombinant Cas9 proteins (250 ng/ml: Thermo Fisher Scientific, MA, USA). The gRNA sequences are as follows:

*Saa1-2* gRNA1: 5′-CCACGUAUGAGGUGGCCCAUGGG-3′

*Saa1-2* gRNA2: 5′-CUGCAGCACACCCACGUAUGAGG-3′

Eggs at the 2-cell stage were transplanted into the oviduct of pseudopregnant mice. F0 mice were crossed with WT and F1 mice were obtained for generating KO mice (≥ F2).

### DNA extraction and genomic PCR

Genomic PCR to genotype mice was performed against DNAs prepared from the mouse tails. The tails were incubated with 50 mM NaOH (nacalai tesque, Kyoto, Japan) for more than 10 min at 95°C. 10 ml of 1 M Tris-HCl pH 8.0 (nacalai tesque) was then added to the reaction, followed by centrifugation at 10,000 × *g* for more than 10 min. The resulting supernatant was subjected to genomic PCR. Genomic PCR for genotyping was performed using KOD FX-Neo (TOYOBO, Osaka, Japan). The primers used in this experiment are shown in Table S1.

### Cell line and cancer transplantation

4T1 mouse breast cancer cell line was cultured and maintained in RPMI1640 (nacalai tesque) in a 5% CO2 tissue culture incubator at 37°C as described previously (Mizuno et al., 2022).

The media (RPMI1640) was supplemented with 10% fetal bovine serum (nacalai tesque) and 1% penicillin/streptomycin (nacalai tesque). The thawed cells were passaged once and then were transplanted to mice. 2.5×10^6^ 4T1 cells resuspended in 100 µL of RPMI1640 containing neither FBS nor penicillin/streptomycin were inoculated subcutaneously into the right flank of an 8–9-week-old BALB/c female mouse. In the sham-treated group, mice were given RPMI1640 supplemented with 10% FBS. Mice were sacrificed on day 14 post-transplantation and the liver and bone marrows were collected.

### RNA isolation, cDNA synthesis, and quantitative reverse transcription PCR (qRT-PCR)

RNA isolation, cDNA synthesis, and qRT-PCR experiments were performed as described previously (Mizuno et al., 2022). Mouse livers were crushed in liquid nitrogen and homogenized with Trizol reagent (Thermo Fisher Scientific, MA, USA). Total RNAs were extracted from the homogenized supernatant using RNeasy Mini Kit (Qiagen, Venlo, Netherlands) according to the manufacturer’s instructions.

The bone marrows were collected essentially as described previously (Amend et al., 2016; Pedersen et al., 2019). Briefly, the mouse right femurs from sham and 4T1-bearing mice were collected, cleaned of muscle tissue, and then were polished with gauze. The femurs were placed with knee-end down in a perforated 0.6 ml tube added 80 μl of RNA*later* (Qiagen) and inserted in a 1.5 ml centrifuge tube, followed by centrifugation at 5,700 × *g* for 30 sec. The resulting pellets were suspended in 1 ml of Trizol reagent (Thermo Fisher Scientific) and proceeded with RNA extraction as described above.

100 to 500 ng of total RNAs were reverse-transcribed with Transcriptor First Strand cDNA synthesis kit (Roche, Basal, Switzerland). qPCR experiments were performed using the StepOnePlus qPCR system (Applied Biosystems, CA, USA) and SYBR Green Master Mix (Roche). *Gapdh* was used as an internal control for the liver samples. *18S rRNA* was used as an internal control for the bone marrow samples.

### Western blotting

Crushed liver powders were lysed with lysis buffer (10 mM Tris-HCl pH8.0, 100 mM KCl, 2.5 mM MgCl_2_, 0.5% TritonX-100, cOmplete (Roche, Basel, Switzerland). The protein concentration was determined using the BCA protein assay kit (TaKaRa, Shiga, Japan) according to the manufacturer’s instructions. 20 μg of the extracted protein were electrophoresed in a 15% SDS polyacrylamide gel for 1 hour at 150V and transferred to a PVDF membrane (Millipore, MA, USA) for 1 hour at 72V. The membrane was incubated with 5% skim milk in Tris-buffered saline, 0.1% Tween20 (TBST) overnight at 4ºC, and then was incubated with mouse serum amyloid A1/A2 antibody (1:1000 in Can Get Signal (TOYOBO, Osaka, Japan): AF2948, R&D Systems, MN, USA) for 2 hours at room temperature. The membrane was washed with TBST three times. Then, the membrane was incubated with goat IgG horseradish peroxidase-conjugated antibody (1:5000 in Can Get Signal (TOYOBO): HAF017, R&D systems) for 1 hour at room temperature. Following TBST-wash steps, signals were visualized using ECL Prime Western Blotting Detection reagent (Cytiva, Tokyo, Japan) and images were taken using Amersham ImageQuant800. The same protein samples were loaded onto a 15% SDS polyacrylamide gel and then stained with SYPRO Ruby Protein Gel Stain (Thermo Fisher Scientific) to confirm an equivalent sample loading in each lane.

### Transcriptome analysis

Total RNAs were extracted as described above with RNase-Free DNase Set (Qiagen). RNA-seq libraries were generated using the NEBNext Globin&rRNA depletion kit and the NEBNext UltraII Directional RNA Library prep kit according to the manufacturer’s instructions (New England Biolabs, MA, USA). Sequencing experiments were performed using NextSeq 500 (Illumina; High Output Kit v2.5, 75 Cycles). The obtained reads were filtered using fastp (version 0.20.1) (Chen et al., 2018) to remove low-quality sequences (< Q30), mapped to the mouse genome (version mm10) using Hisat2 (version 2.2.1) (Kim et al., 2019), and processed using Samtools (version 1.10) (Danecek et al., 2021) and featureCounts (version 2.0.1) (Liao et al., 2014). Read counts were normalized with the reads per million per kilobase (RPKM) method. The generated gene expression matrix with RPKM scores is listed in Table S2.

Gene expression matrix was used to perform gene set enrichment analysis (GSEA) to interpret transcriptional profiles (Subramanian et al., 2005; Wang et al., 2013). Enrichment score (ES), normalized enrichment score (NES), and false discovery rate (FDR) for all variables and signatures were obtained. The differentially expressed genes (DEGs) were defined as genes with |log_2_(fold change)| >1 and *p* value < 0.05. The volcano plots were depicted using ggplot2 to visualize DEGs (https://ggplot2.tidyverse.org/index.html). ImmuCellAI-mouse (Miao et al., 2021; Miao et al., 2020), a bulk RNA-seq data deconvolution approach, was applied to estimate the abundance of 36 immune cell types using the default settings.

### Flow cytometry

The livers of sham and 4T1-bearing mice were harvested on day 14 after transplantation. The obtained liver tissues were homogenized in 8 mL of RPMI1640 media containing 1.8 mg/ml Collagenase IV (WOR-CLS4-1, Worthington Biochemical Corporation, NJ, USA) and 112.3 µg/ml DNase I (11284932001, Roche). 8 ml of the suspension was filtered using a cell strainer (70 µm mesh) and then mixed with 4 ml of 90 % Percoll solution (Sigma-Aldrich, MO, USA). The suspension was then centrifuged at 700 × *g* for 20 min. The red blood cells in the pellets were lysed with 1×Lysing buffer (Lysing Buffer 10× Concentrate (BD Biosciences) diluted by H_2_O (nacalai tesque)). The samples were washed with RPMI1640 media supplemented with 2% FBS. The obtained samples were stained for 15 min on ice in FACS buffer (2% FBS, 0.05% NaN_3,_ and 1×PBS) mixed with TruStain FcX™ (anti-mouse CD16/32) Antibody (1:200, Clone: 93, BioLegend, CA, USA) and eBioscience™ Fixable Viability Dye eFluor™ 780 (1:1000, Invitrogen, MA, USA). The samples were washed with 100 μl of FACS buffer. The washed samples were stained for 20 min on ice with Brilliant Violet 510™ anti-mouse CD45 Antibody (1:200, Clone: 30-F11, BioLegend), PE/Cyanine7 anti-mouse/human CD11b Antibody (1:200, Clone: M1/70, BioLegend), PE anti-mouse Ly-6C Antibody (1:200, Clone: HK1.4, BioLegend), and FITC anti-mouse Ly-6G Antibody (1:200, Clone: 1A8, BioLegend), and APC anti-mouse F4/80 Antibody (1:50, Clone: BM8, BioLegend) in FACS buffer. Following a wash step using FACS buffer, the stained samples were resuspended with FACS buffer and then filtered using a cell strainer (35 μm mesh) set in a 5 mL tube (Falcon). The resulting samples were analyzed using FACS Canto II (BD Bioscience, NJ, USA) and calculated using FlowJo software (v10.801) (BD Biosciences).

### Statistics and data visualization

GraphPad Prism Software was used to analyze data. Data were displayed as mean ± SEM. Student’s *t* test was performed to analyze the statistical significance between groups unless otherwise indicated, and *p* value < 0.05 was considered statistically significant.

### Data availability

RNA-seq data obtained in this study are available from DNA Databank of Japan under the accession number of DRA014884.

## Acknowledgement

This work was supported by JSPS KAKENHI (17H06299, 18K15409, 18H04810, 20H03451, and 20H04842; S.K), JST FOREST (20351876; S.K), JST Moonshot (JPMJMS2011-61; S.K), Caravel, Co., Ltd (S.K), Ono Medical Research Foundation (S.K), Takeda Science Foundation, The Uehara memorial foundation (S.K), Chubei Ito foundation (S.K), Japan Foundation for applied Enzymology (S.K) and the scholarship from China Scholarship Council under the Grant 202008050199 (C.H).

We thank Hitoshi Miyachi and Satsuki Kitano for their help in the generation of *Saa1-2* KO mice. We thank Dr. Takefumi Kondo and Yukari Sando for their support in the transcriptome analyses. We also thank Daiya Ohara and Dr. Keiji Hirota for their advices on the neutrophil-related experiments.

## Competing interests

The authors declare no competing interests exist in this study.

## Author contribution

C.H and R.K performed experiments, analyzed data, constructed figures. A.H, Y.N, R.M, M.Y performed experiments. M.T substantially contributed to the conception of this study. KK supervised the study. SK conceived and supervised the study, analyzed data, and wrote the paper. All authors provided intellectual input and reviewed the paper.

---

**Figure S1:**
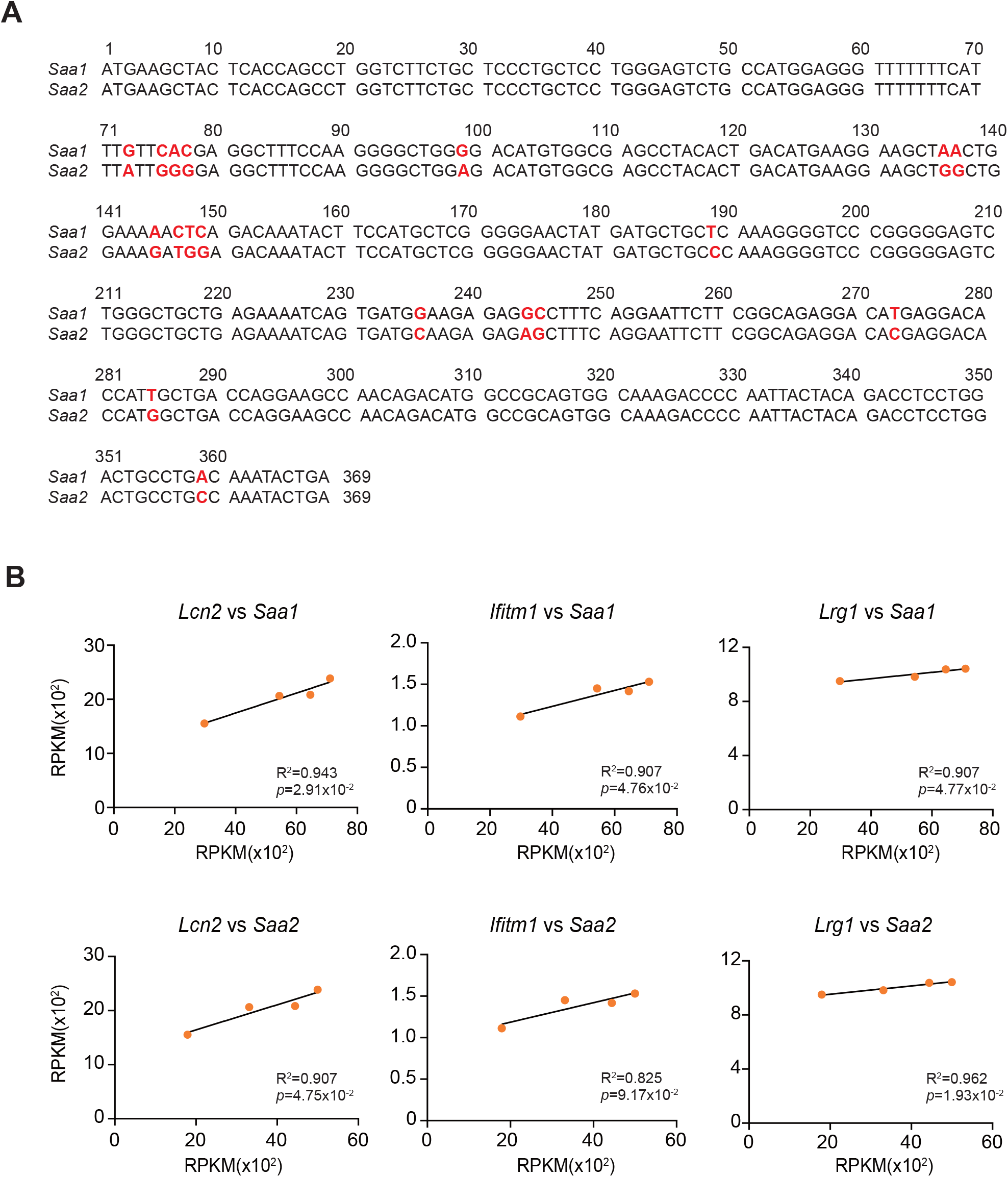
The basic characteristics of *Saa1-2*. **(a)** A DNA sequence alignment of *Saa1* (accession number: CCDS21284.1) and *Saa2* (accession number: CCDS21285.1). The differences in their sequences are highlighted in red. **(b)** The correlations of the mRNA abundances between *Saa1-2* and representative neutrophil-related genes. *n* = 4.

**Figure S2:**
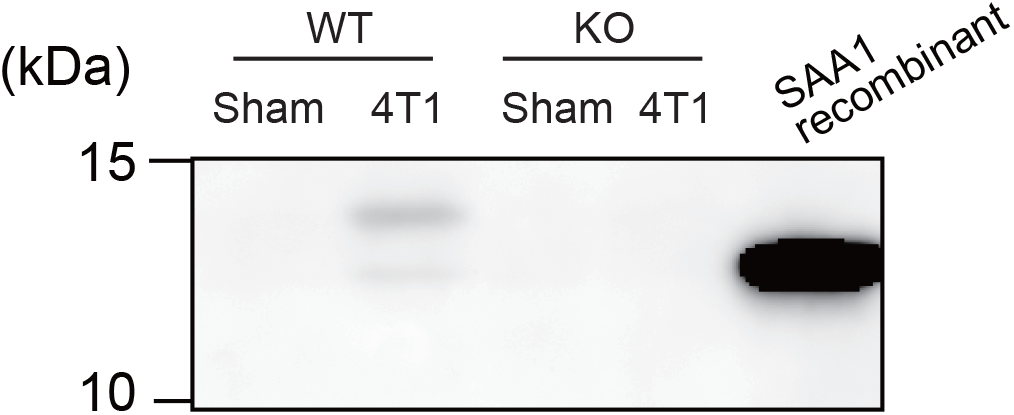
Western blot analysis of SAA1-2 proteins in the liver. Western blot analysis for SAA1-2 in the livers of sham and 4T1-bearing mice in WT and *Saa1-2* KO. Recombinant SAA1 protein (2948-SA: R&D systems, MN, USA) is included as a positive control.

**Figure S3:**
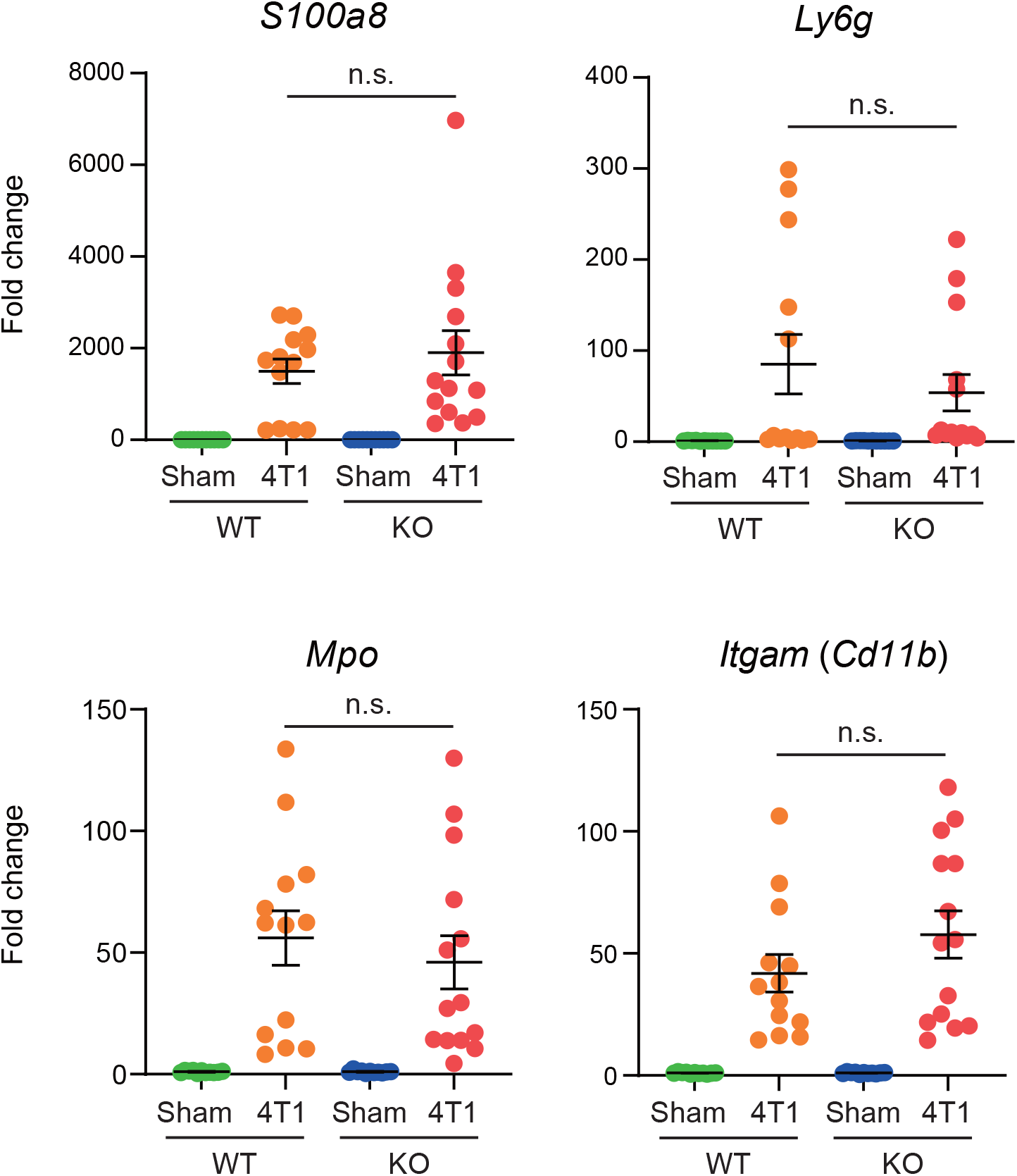
qPCR validations of the RNA-seq experiments in the liver. qPCR analysis of *S100a8, Ly6g, Mpo*, and *Itgam* (*Cd11b*) in the livers of sham and 4T1-bearing mice in WT and *Saa1-2* KO (pooled from the four independent 4T1 transplantation experiments). Averaged fold change data normalized to the sham group in each genotype are presented as the mean ± SEM. *n.s*., not significant, unpaired two-tailed Student’s *t*-test. *n* = 11 for the sham groups, *n* = 13 for 4T1-bearing WT mice, and *n* = 14 for 4T1-bearing *Saa1-2* KO mice.

**Figure S4:**
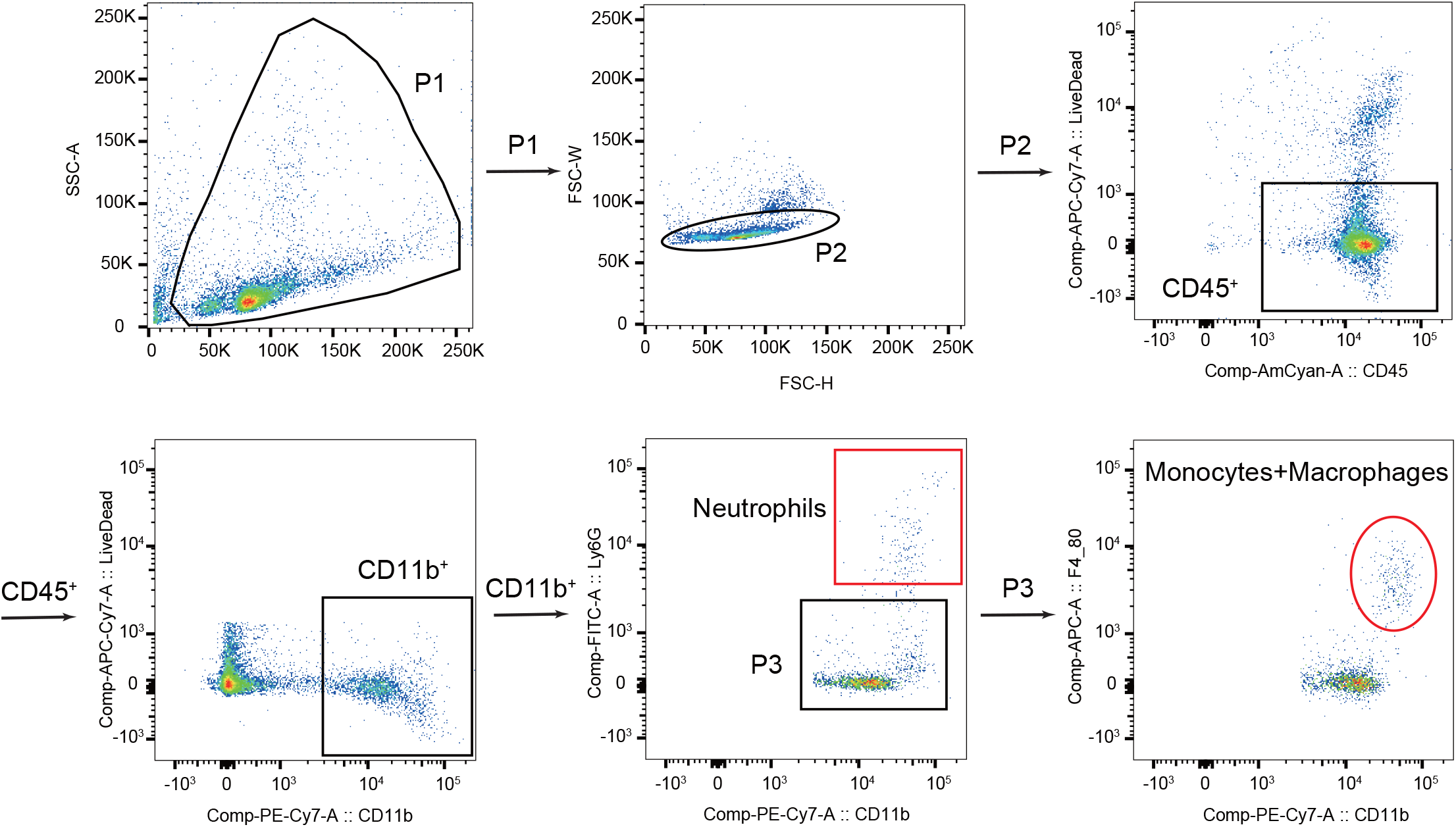
Gating strategies in the flow cytometry experiments.

**Figure S5:**
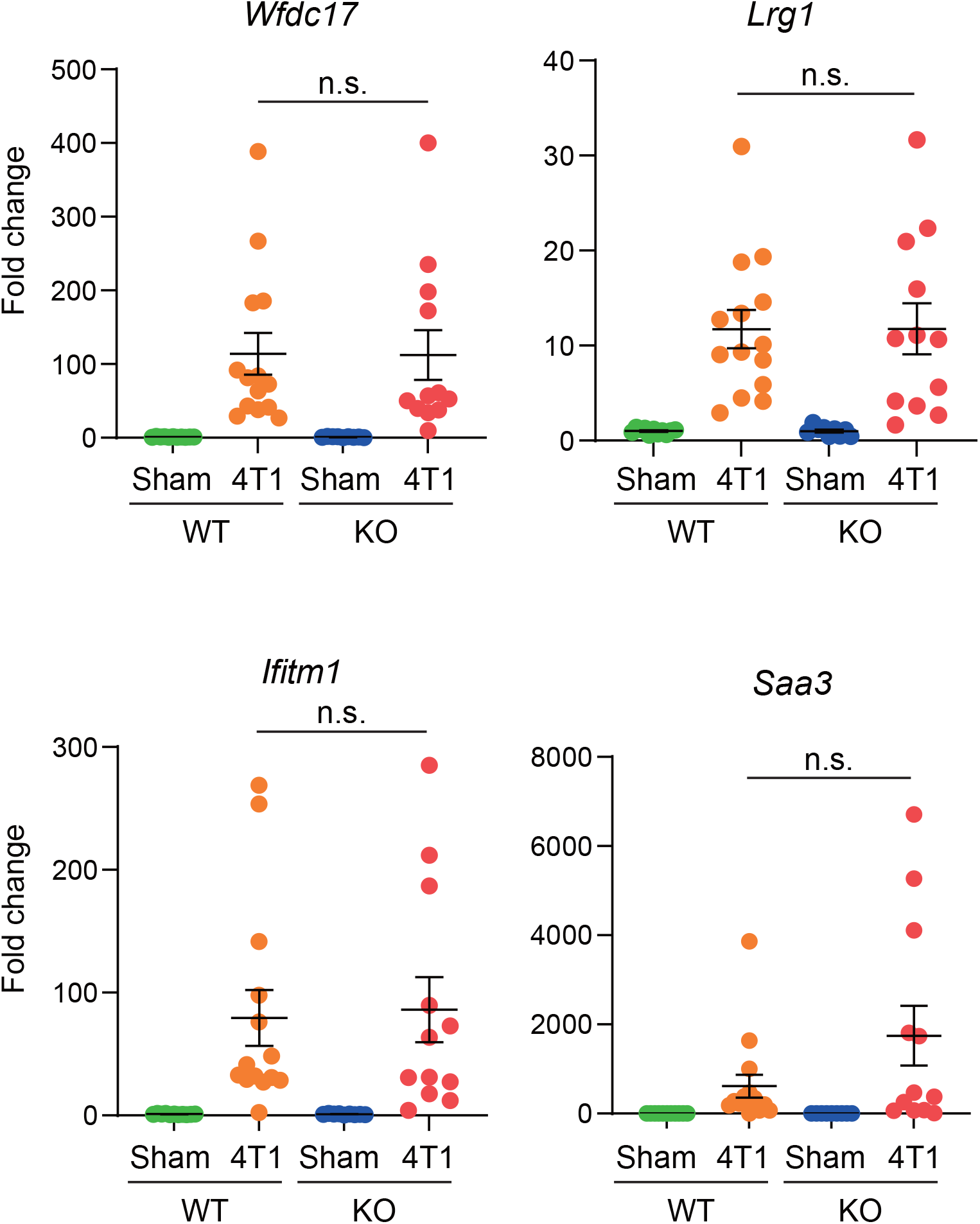
qPCR validations of the RNA-seq experiments in the bone marrow. qPCR analysis of *Wfdc17, Lrg1, Ifitm1*, and *Saa3* in the bone marrows of sham and 4T1-bearing mice in WT and *Saa1-2* KO (pooled from the four independent 4T1 transplantation experiments). Data are normalized with *18s rRNA* and are presented as the mean ± SEM. *n.s*., not significant, unpaired two-tailed Student’s *t*-test. *n* = 11 for sham-treated WT mice, *n* = 14 for 4T1-bearing WT mice, *n* = 9 for sham-treated WT mice, and *n* = 12 for 4T1-bearing *Saa1-2* KO mice.

**Figure S6:**
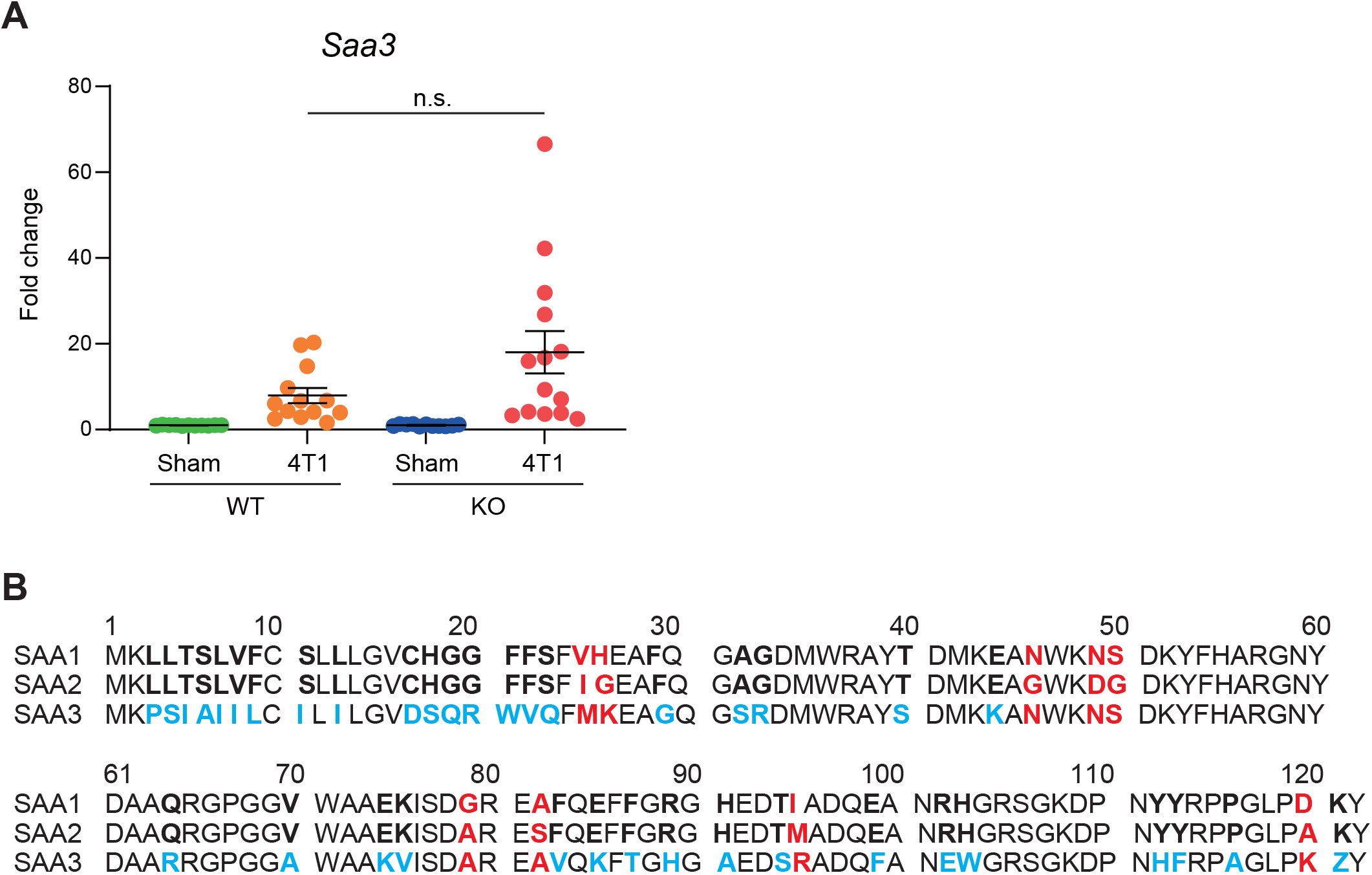
*Saa3* is induced in the livers of 4T1-bearing mice even in the absence of *Saa1-2*. **(a)** qPCR analysis of *Saa3* in the livers of sham and 4T1-bearing mice in WT and *Saa1-2* KO (pooled from the four independent 4T1 transplantation experiments). Averaged fold change data normalized to the sham group in each genotype are presented as the mean ± SEM. *n.s*., not significant, unpaired two-tailed Student’s *t*-test. *n* = 11 for the sham groups, *n* = 13 for 4T1-bearing WT mice, and *n* = 14 for 4T1-bearing *Saa1-2* KO mice. **(b)** An amino acid sequence alignment of SAA1 (accession number: CCDS21284.1), SAA2 (accession number: CCDS21285.1) and SAA3 (accession number: CCDS21282.1).

## Notes

### Competing Interest Statement

The authors have declared no competing interest.

